# Cross-talk between cAMP and Ca^2+^ signalling in cardiac pacemaker cells involves IP_3_-evoked Ca^2+^ release and stimulation of adenylyl cyclase 1

**DOI:** 10.1101/2022.01.06.475211

**Authors:** Samuel J. Bose, Matthew Read, Rebecca A Capel, Emily Akerman, Thamali Ayagama, Angela Russell, Derek A Terrar, Manuela Zaccolo, Rebecca AB Burton

**Author notes:** **Correspondence to:** Dr Rebecca-Ann B Burton, Department of Pharmacology, University of Oxford, Mansfield Road, Oxford, Oxon, OX1 3QT.

## Abstract

Atrial arrhythmias, such as atrial fibrillation (AF), are a major mortality risk and a leading cause of stroke. The IP_3_ signalling pathway has been proposed as an atrial specific target for AF therapy, and atrial IP_3_ signalling has been linked to the activation of calcium sensitive adenylyl cyclases AC1 and AC8. Here we investigated the involvement of AC1 in the response of intact mouse atrial tissue and isolated guinea pig atrial and sinoatrial node (SAN) cells to the α-adrenoceptor agonist phenylephrine (PE) using the selective AC1 inhibitor ST034307. The maximum rate change of spontaneously beating mouse right atrial tissue exposed to PE was reduced from 14.46 % to 8.17% (*P* = 0.005) in the presence of 1 μM ST034307, whereas the increase in tension generated in paced left atrial tissue in the presence of PE was not inhibited by ST034307 (Control = 14.20 %, ST034307 = 16.32 %; *P* > 0.05). Experiments were performed using isolated guinea pig atrial and SAN cells loaded with Fluo-5F-AM to record changes in calcium transient amplitude (CaT) generated by 10μM PE in the presence and absence of 1μM ST034307. ST034307 significantly reduced the beating rate of SAN cells (0.34-fold decrease; *P* = 0.004), but did not result in an inhibition of CaT amplitude increase in response to PE in atrial cells. The results presented here demonstrate the involvement of AC1 in the downstream response of atrial pacemaker activity to α-adrenoreceptor stimulation and IP_3_R calcium release.

## Introduction

Effective Ca^2+^ handling is essential to the normal physiological function of the heart (Bers, 2002). Cardiac activity is closely regulated by the action of Ca^2+^ dependent enzymes, such as calcineurin and Ca^2+^/calmodulin-dependent kinase II (CaMKII), as well as Ca^2+^ mobilising agents such as inositol-1,4,5-trisphosphate (IP_3_), cyclic ADP-ribose (cADPR) and nicotinic acid adenine-dinucleotide phosphate (NAADP) (Terrar, 2020). In addition to the regulation of excitation-contraction coupling (ECC), the process by which electrical activity is converted into cardiac contraction, currents that contribute to the pacemaker potential in the sinoatrial node (SAN) and atrioventricular node (AVN) are highly dependent on the regulation of intracellular Ca^2+^ signalling (Hancox & Mitcheson, 1997; Lakatta *et al*., 2010; Capel & Terrar, 2015; Burton & Terrar, 2021).

### cAMP signalling in the heart

Under normal physiological conditions, both cardiac ECC and pacemaker activity are primarily regulated via the combination of β-adrenergic and muscarinic signalling, leading to downstream activation of adenylyl cyclase, and subsequent generation of cyclic-adenosine monophosphate (cAMP) (Bers, 2002; Burton & Terrar 2021). The predominant AC isoforms expressed in ventricular myocytes are AC5 and AC6 (Katsushika *et al*., 1992; Premont *et al*., 1992). Traditionally, cAMP has been thought to act primarily via protein kinase A (PKA) (Walsh *et al*., 1968; Krebs & Bevo, 1979), and cyclic nucleotide-gated ion channels (Fesenko *et al*. 1985) to influence cardiomyocyte contractile sensitivity as well as regulating L-type Ca^2+^ channel (LTCC) activity (Chen-Izu *et al*., 2000; Harvey & Clancy, 2021). However more recent work has demonstrated that cAMP may also act via exchange proteins directly activated by cAMP (EPACs) (de Rooij *et al*., 1998), as well as ‘Popeye domain’ containing proteins (Brand *et al*. 2005; Zaccolo *et al*. 2021). The downstream effects of cAMP signalling in cardiomyocytes can therefore influence a wide range of cellular processes, including gene expression and cell morphology, in addition to electrical and contractile activity (Harvey & Clancy, 2021). This heterogeneity in function is the result of spatial confinement and localized compartmentalisation of cAMP signalling within subcellular nanodomains (Zaccolo & Pozzan, 2002; Zaccolo *et al*. 2021).

### Role of calcium-sensitive adenylyl cyclases in the atria

In addition to the role of AC5 and AC6 in response to β-adrenergic signalling, there is also a role for α-adrenergic signalling, and resultant IP_3_ production, in the regulation of cardiac activity (Domeier *et al*., 2008; Terrar, 2020; Burton & Terrar, 2021; Capel *et al*., 2021). Increasing evidence supports a mechanism whereby activation of the IP_3_ signalling pathway acts through the downstream activation of the Ca^2+^ activated adenylyl cyclases AC1 and AC8 (Georget *et al*., 2002; Mattick *et al*., 2007; Burton & Terrar, 2021; Capel *et al*., 2021). In guinea pig atrial cardiomyocytes, AC1 and AC8 are localised in the plasma membrane in close proximity to type 2 IP_3_ receptors (IP_3_R2) on the sarcoplasmic reticulum (SR) (Capel *et al*., 2021). Inhibition of ACs using MDL-12,330A, or inhibition of PKA using H89, inhibits the increase in Ca^2+^ transient amplitude observed in isolated guinea pig atrial cardiomyocytes in response to either intracellular photorelease of caged IP_3_ (IP_3_/PM) or external stimulation of α-adrenergic signalling using phenylephrine (PE), thereby demonstrating downstream activation of AC’s by the IP_3_ signalling pathway.

### Role of calcium-sensitive adenylyl cyclases in the SAN

In the SAN, spontaneous pacemaker activity is the result of the ‘coupled-clock’ mechanism, involving tight coupling between rhythmic Ca^2+^ release from the SR (i.e. ‘Ca^2+^ clock’), and rhythmic oscillations in the membrane potential (i.e. ‘membrane clock’) (Lakatta *et al*., 2010; Tsutsui *et al*., 2018). Indeed, it appears this coupling is essential for pacemaking in human SAN cells (Tsutsui *et al*., 2018). Membrane clock activity results from the alternation and balance between depolarising currents (e.g. I_f_, I_CaL_, I_sust_) and repolarising currents (e.g. I_Ks_ and I_Kr_) (Difrancesco & Tromba, 1988; Difrancesco *et al*., 1991; Burton & Terrar, 2021). The hyperpolarisation activated ‘funny’ current I_f_, is carried by the HCN channel, and modulated by changes in cytosolic Ca^2+^ (Rigg *et al*., 2003), as well as sub-sarcolemmal cAMP (Difrancesco & Tortora, 1991; Burton & Terrar, 2021). I_f_, and therefore SAN pacemaking, can thereby be influenced by phosphodiesterase (PDE) and AC activity (Difrancesco & Tortora, 1991; Mattick *et al*., 2007; Vinogradova *et al*., 2008).

The increases in spontaneous beating rate observed in intact murine right atria in response to PE is similarly inhibited using either MDL-12,330A or H89 (Capel *et al*., 2021), suggesting a role for Ca^2+^-activated AC1 and or AC8 in the positive inotropic and chronotropic response to IP_3_ signalling in cardiac atria. AC1 activity modulates the I_f_ current in the SAN in the absence of β-adrenergic stimulation, and contributes to the higher resting cAMP level in SAN cells compared to ventricular cells (Mattick *et al*., 2007). These observations suggest cAMP signalling, downstream of AC1 activation, is a crucial mechanism by which the Ca^2+^ clock and membrane clock are coupled in the SAN. Furthermore, the modulation of I_f_ by cytosolic Ca^2+^ appears to be independent of CaMKII as chelation of SAN Ca^2+^ using BAPTA reduces I_f_, whereas inhibition of CaMKII is without effect (Rigg *et al*., 2003). Although CaMKII is essential for SAN pacemaker activity (Yaniv *et al*., 2010), the actions of CaMKII on pacemaker function are linked to effects on I_Ca,L_ rather than I_f_ (Vinogradova *et al*. 2000; Rigg *et al*. 2003). Conversely, inhibition of SAN ACs using the non-specific AC inhibitor MDL-12,330A reduces I_f_, whilst inhibition of phosphodiesterase using IBMX to inhibit the breakdown of basal cAMP increases I_f_, suggesting a role for Ca^2+^-activated ACs in regulating the I_f_ current in SAN cells (Mattick *et al*., 2007).

In the present study, we aimed to investigate the role of AC1 in both atrial and SAN IP_3_ signalling using the chromone-derivative ST034307 (originally identified by chemical library screening), a small molecule AC1 inhibitor that is selective for AC1 over other AC isoforms, including ACs 2,5,6 and 8, at concentrations below 30 μM (IC_50_ = 2.3 μM) (Brust *et al*., 2017). Specifically, the role of AC1 in IP_3_-mediated potentiation of SAN pacemaker activity and atrial inotropy was investigated by using mouse atrial preparations as well as isolated guinea-pig SAN and atrial cells.

## Materials and Methods

### Animals

All animal experiments were performed in accordance with the United Kingdom Home Office Guide on the Operation of Animal (Scientific Procedures) Act of 1986. All experimental protocols (Schedule 1) were approved by the University of Oxford, Procedures Establishment License (PEL) Number XEC303F12.

### Drugs and reagents

AC1 was inhibited using the AC1 selective inhibitor ST034307 (Tocris, UK) (Brust *et al*., 2017). In all experiments, ST034307 was dissolved in DMSO to make 3mM stock prior to addition to experimental solutions at a final concentration of 1 μM and applied for at least 5 min for isolated cells, or 30 min for whole-tissue experiments in order to ensure sufficient tissue penetration.

### Atrial myocyte isolation

Male Dunkin Hartley guinea pigs (300-550g, Envigo, UK) were housed and maintained in a 12 h light-dark cycle with *ad libitum* access to standard diet and sterilized water. Guinea pigs were culled by concussion followed by cervical dislocation in accordance with Home Office Guidance on the Animals (Scientific Procedures) Act (1986). Atrial myocytes were isolated following the method of Collins *et al*. (2011). Hearts were rapidly excised and washed in physiological salt solution (PSS, in mM): NaCl 125, NaHCO_3_ 25, KCl 5.4, NaH_2_PO_4_ 1.2, MgCl_2_ 1, glucose 5.5, CaCl_2_ 1.8, oxygenated with 95 % O_2_ / 5 % CO_2_ (solution pH 7.4 after oxygenation and heating) to which heparin was added (final concentration = 20 IU·ml^-1^) to prevent clot formation in the coronary vessels. Hearts were then mounted on a Langendorff apparatus for retrograde perfusion via the aorta. Perfusion was initially carried out in a modified Tyrode solution containing (mM): NaCl 136, KCl 5.4, NaHCO_3_ 12, Na^+^ pyruvate 1, NaH_2_PO_4_ 1, MgCl_2_ 1, EGTA 0.04, glucose 5; gassed with 95% O_2_/5% CO_2_ to maintain a pH of 7.4 at 35+/-1°C. After 2 min this was replaced with a digestion solution: the modified Tyrode above containing 100 μM CaCl_2_ and either 0.6 mg/mL of collagenase (type II, Worthington Biochemical Corp., Lakewood, NJ, USA) or 0.02-0.04 mg/ml Liberase™ TH (Roche, Penzberg, Germany), but no EGTA.

After this enzymatic digestion, the heart was removed from the cannula and the atria were separated from the ventricles. For the isolation of atrial myocytes, slices of the atria were triturated using a glass pipette) and stored at 4°C in a high potassium medium containing (in mM): KCl 70, MgCl_2_ 5, K^+^ glutamine 5, taurine 20, EGTA 0.1, succinic acid 5, KH_2_PO_4_ 20, HEPES 5, glucose 10; pH to 7.2 with KOH. For the isolation of SAN cells, the translucent SAN region, located on the upper surface of the right atrium, in between the inferior and superior vena cava (Rigg *et al*., 2003), immediately medial to the crista terminalis was identified under a dissection microscope. Thin tissue strips encompassing the nodal region were dissected, triturated using a glass pipette and stored at 4°C in high potassium medium. For experiments, healthy atrial myocytes were identified based on morphology, and healthy SAN myocytes by morphology and the presence of spontaneous, rhythmic beating in the absence of electrical stimulation.

### Immunocytochemistry

Immunocytochemical labelling and analysis was carried out using the method of Collins and Terrar (2012). Rabbit anti-AC1 (55067-1-AP) primary antibody was purchased commercially (ProteinTech, Manchester, United Kingdom) and used at a dilution of 1:200. Mouse anti-IP_3_R monoclonal primary antibody IP_3_R2 (sc-398434) was purchased commercially (Santa Cruz Biotechnology, Santa Cruz, CA, USA) and used at a dilution of 1:100. The specificity of antibody sc-398434 has been previously verified using Western blot by Lou et al. (2021). IP_3_R antibodies have been extensively covered in previous studies (Hattori *et al*., 2004; Ando *et al*., 2006; Uchida *et al*., 2010; Salvador & Egger, 2018). Isolated cardiac cells were plated onto flamed coverslips and left to adhere for 30 min at 4°C. Cells were first fixed in 4% paraformaldehyde–phosphate buffered saline (PBS) for 15 min. Once the cells were fixed, they were washed in PBS (3 changes, 5 min each). Cells were then permeabilised and blocked using the detergent Triton X-100 (0.1%), 10% normal donkey and 10% BSA in PBS (Sigma-Aldrich) for 60 min at room temperature to reduce non-specific binding. After blocking, the cells were incubated with primary antibodies at 4°C overnight dissolved in blocking solution. The next day, cells were first washed with PBS (3 changes, 5 min each) before being incubated with secondary antibodies; AlexaFluor -488 or -546 conjugated secondary antibodies (Invitrogen, UK), raised against the appropriate species, at RT for 120 min in PBS then washed with PBS (3 changes, 5 min each). Finally, the cells were mounted using Vectashield with DAPI and permanently sealed with nail polish.

Cells were stored in the dark at 4°C and imaged within 2 days. Observations were carried out using an AXIO Observer Z1 confocal microscope (LSM 710; Carl Zeiss) with a 63x/1.4 numerical aperture Plan-Apochromat oil objective lens. ZEN Black 2008 (Zeiss) was used to acquire multichannel fluorescence images. For detection of AlexaFluor 488, fluorescence excitation was at 488 nm (argon laser) with emission collected at 505-530 nm. Excitation at 543 nm (HeNe laser) was collected at >560 nm for detection of AlexaFluor 546. The two channels were imaged sequentially at 8-bit. Z-stack images were collected at 1 μm sections.

### Murine atrial studies

Adult male CD1 mice (30-35 g, Charles River, UK) were housed in a 12 h light-dark cycle with *ad libitum* access to standard diet and sterilized water. Mice were culled by concussion followed by cervical dislocation in accordance with Home Office Guidance on the Animals (Scientific Procedures) Act (1986). The heart was rapidly excised and washed in heparin-containing PSS. The ventricles were dissected away under a microscope and the atria were cleared of connective tissue before being separated. Right and left atrial preparations were mounted separately in a 37 °C organ bath containing oxygenated PSS and connected to a force transducer (MLT0201 series, ADInstruments, UK) in order to visualize contractions. Resting tension was set between 0.2 and 0.3 g, and the tension signals were low-pass filtered (20 Hz for right atria and 25 Hz for left atria). Right atrial beating rate was calculated from the time interval between contractions. Left atria were electrically field stimulated at a constant rate of 5 Hz using a custom-built stimulator connected to coil electrodes positioned both sides lateral to the left atrial tissue. Voltage was set at the threshold for stimulating contraction plus 5 V, and was within the range 10-20 V or all experiments. In all experiments, preparations were allowed to stabilise at a resting beating rate (>300 bpm) in PSS for 30 minutes. After stabilisation (variation in average rate of a 10s sample of no more than 2 bpm over a 10-minute period or 0.01 g change in tension), metoprolol (1 μM) was added to the bath to ensure specificity to α-adrenergic effects, plus or minus ST034307 (1 μM). Each addition was allowed to stabilise for a further 30 minutes or until stability was achieved as described above. Cumulative concentrations of PE were added to the bath at intervals of 5 minutes (range 0.1 to 30 μM) in the presence of metoprolol. Preparations were excluded if stabilized beating rate under control conditions (PSS only) was less than 300 bpm, in the case of the right atrium, or if preparations were not rhythmic. Data were fitted using Log(agonist) versus response curves (three parameter model) by nonlinear regression using a least squares method (Prism v9). AC1 was inhibited using the AC1 selective inhibitor ST034307 (Tocris, UK) (Brust *et al*., 2017). ST034307 (1 μM) was added after stabilization of the preparations in the presence of metoprolol and applied for at least 30 min prior to PE additions. PE dose-response curves were started only after tissue had reached a stable response.

### Ca^2+^ transient imaging

For whole-cell fluorescence experiments, isolated atrial myocytes were incubated with membrane permeant Fluo-5F-AM (3 μM) for 10 min then plated to a glass cover slip for imaging. Cells were incubated for a further 10 minutes in-situ in the organ bath to allow time for cells to adhere to the cover slip before perfusion with PSS. Carbon fibre electrodes were used to field-stimulate Ca^2+^ transients at a rate of 1 Hz. All experiments were carried out at 35 ± 2°C (fluctuation within a single experiment was <0.5°C) under gravity-fed superfusion with PSS, oxygenated with 95 % O_2_ / 5 % CO_2_ (solution pH 7.4 after oxygenation and heating). Solution flow rate was 3 mL min^-1^. Cells were visualized using a Zeiss Axiovert 200 with attached Nipkow spinning disc confocal unit (CSU-10, Yokogawa Electric Corporation, Japan). Excitation light, transmitted through the CSU-10, was provided by a 488 nm diode laser (Vortran Laser Technology Inc., Sacramento, CA, USA). Emitted light was passed through the CSU-10 and collected by an Andor iXON897 EM-CCD camera (Oxford Instruments, UK) recorded at a minimum acquisition frame rate of 60 frames per second using μManager software (v2.0) and imageJ (Exposure time = 3-10 ms; binning = 4×4). In order to avoid dye bleaching the cells were not continually exposed to 488 nm light. Instead, a video of 8-10 s of Ca^2+^ transients was recorded at discrete timepoints. Ca^2+^ transient time courses were analysed in imageJ and ClampFit (version 10.4). For analysis of Ca^2+^ transient rise and decay times Ca^2+^ data were analysed using pClamp v10 (Molecular Devices, CA, USA) to generate times corresponding to 10-90% and 10-50% rise time, and 90-10%, 90-75%, 90-50% and half width decay time. Decay phases of transients were also fitted using one phase decay least squares regression (Prism v9).

### Statistics

Data were tested for normality by using a Shapiro-Wilk test in Prism v9 software (GraphPad, CA, USA). For all single cell data, two-way paired t-tests, repeated measures 2-way ANOVA or mixed effects analysis were used as appropriate, with Dunnett’s or Tukey’s post hoc test to compare groups to a single control or to all other groups as required (alpha = 0.05). For SAN data, Sidak’s multiple comparison was used to compare control and ST034307 data at all time points. Log[concentration]-response curves, used to estimate EC50s and maximum responses, were calculated using Prism v9 software (GraphPad, CA, USA), by fitting an agonist-response curve with a fixed slope to normalized response data. Normalized data was used to compare responses. Fitted values were compared using 2-way repeated measures ANOVA followed by Šídák’s multiple comparisons or Fisher;s LSD test. Data are presented as mean ± SEM of recorded values, other than dose-response curve maximums and EC50 which are given as mean ± 95 % confidence interval of best-fit value.

## Results

### Immunohistochemistry suggests IP_3_R2 and AC1 are colocalised in isolated guinea pig atrial myocytes

To determine structural and anatomical characterisation of AC1 and IP_3_R2 in atrial myocytes from healthy guinea pig adult hearts, isolated right atrial myocytes were fixed and immunolabelled for AC1 and IP_3_R. Figure 1 shows representative confocal images of atrial myocytes stained with primary antibodies raised against the AC1 (cyan) and IP3R (magenta) proteins. These results along with previous data from our group (Capel *et al*., 2021) showed IP_3_R2 were present on the sarcoplasmic reticulum membrane in atrial myocytes (Figure 1Ai) and AC1 puncta are located in close proximity to IP_3_R2 (Figure 1Aiv, pixel size = 0.264 × 0.264 μm), suggesting Ca^2+^ release from IP_3_R could contribute to the Ca^2+^-induced calcium release process. ImageJ intensity analysis revealed, pixel by pixel by line intensity plots (Figure 1A and B) and in whole image intensity plots (Figure 1C-D), levels of colocalization between AC1 and IP_3_R2 in isolated guinea pig atrial myocytes comparable to that reported by Capel *et al* (2021) (*R* = 0.48 ± 0.05 *n* = 5). Z-stack images showing cross sections through the cell at intervals of 1 μm have been presented in Supplementary Video 1. These results together with recently published data (Capel *et al*., 2021), suggest that IP_3_R-dependent signalling may be capable of stimulating Ca^2+^-dependent ACs and the close positioning of AC1 to IP_3_R2 suggests that AC1 may be an effector of this interaction.

**Figure 1.**
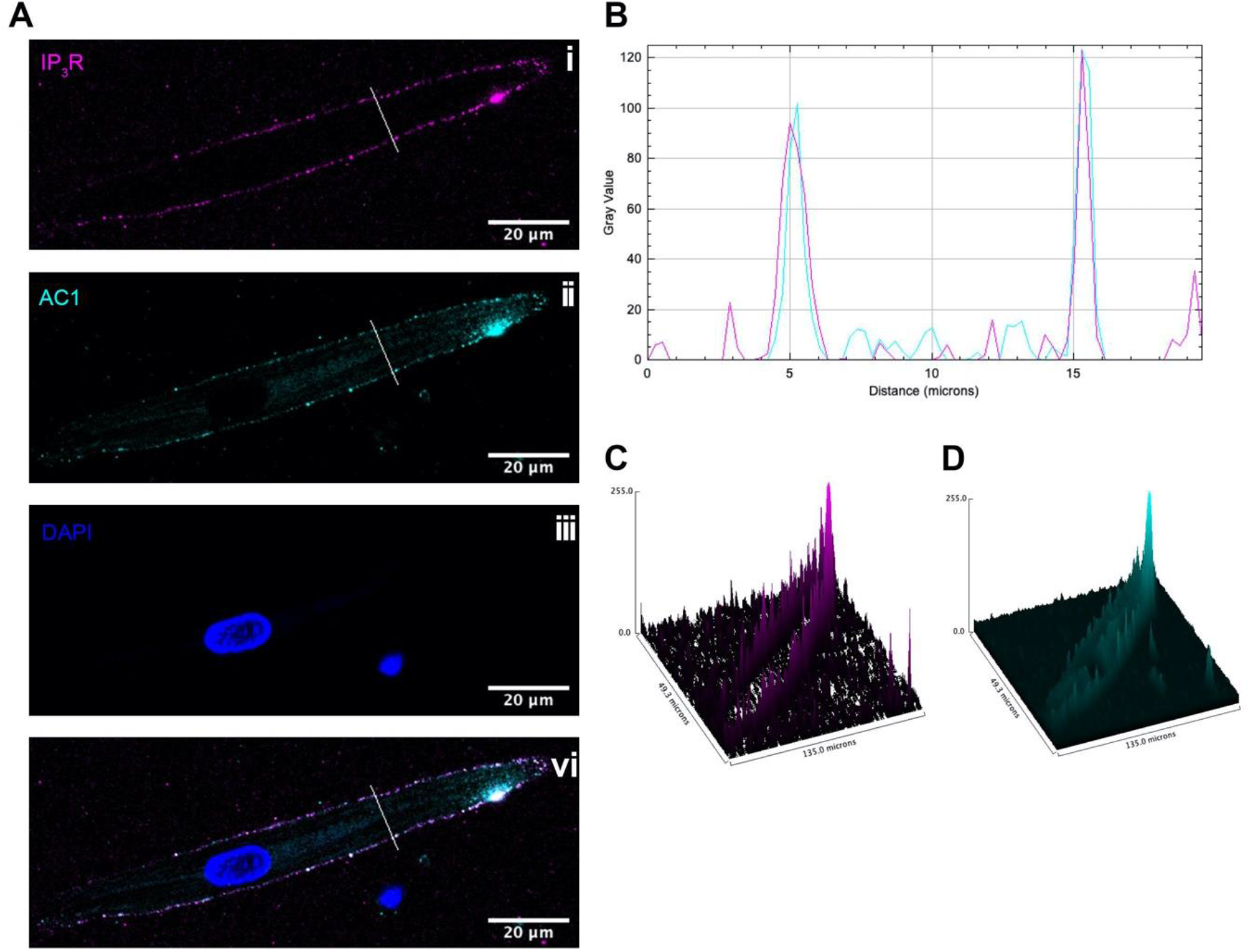
IP_3_R2 is expressed in close proximity with AC1 in guinea pig atrial myocytes. **A**: Representative example of a fixed, isolated guinea pig atrial myocyte (**i**) co-immunolabelled for IP_3_R2 (magenta), (**ii**) AC1 (cyan), (**iii**) DAPI and (**iv**) co-immunolabelled for IP_3_R2 (magenta) and AC1 (cyan). **B**: Intensity plot to show staining intensity along the line shown in **A. C-D**: Intensity surface plot showing the distribution of staining of IP_3_R2 (magenta, **C**) and AC1 (cyan, **D**) for the whole cell, as shown in **Aiii**. Scale bars representing 20 μm are indicated in **A**. For the purposes of presentation only, red and green channels have been represented as magenta and cyan respectively and the intensity of the magenta signal has been amplified by a factor of 8.

### Inhibition of AC1 by ST034307 reduces the positive chronotropic effect of PE in spontaneously beating right atria

Intact, spontaneously beating right atrial tissue preparations can be used to indirectly record SAN activity through the measurement of beating rate, whilst intact left atria can be used to record changes in contractile force generated when stimulated at a constant rate and voltage (Capel *et al*., 2021). Isolated murine right and left atria were mounted separately in organ baths and perfused with PSS at 37 °C in the presence of 1.0 μM metoprolol to inhibit β-adrenergic signalling. Dose response curves for either spontaneous beating rate (right atria, Figure 2A) or tension generated (left atria, Figure 2B) were generated in response to 0.1 to 30 μM PE.

**Figure 2.**
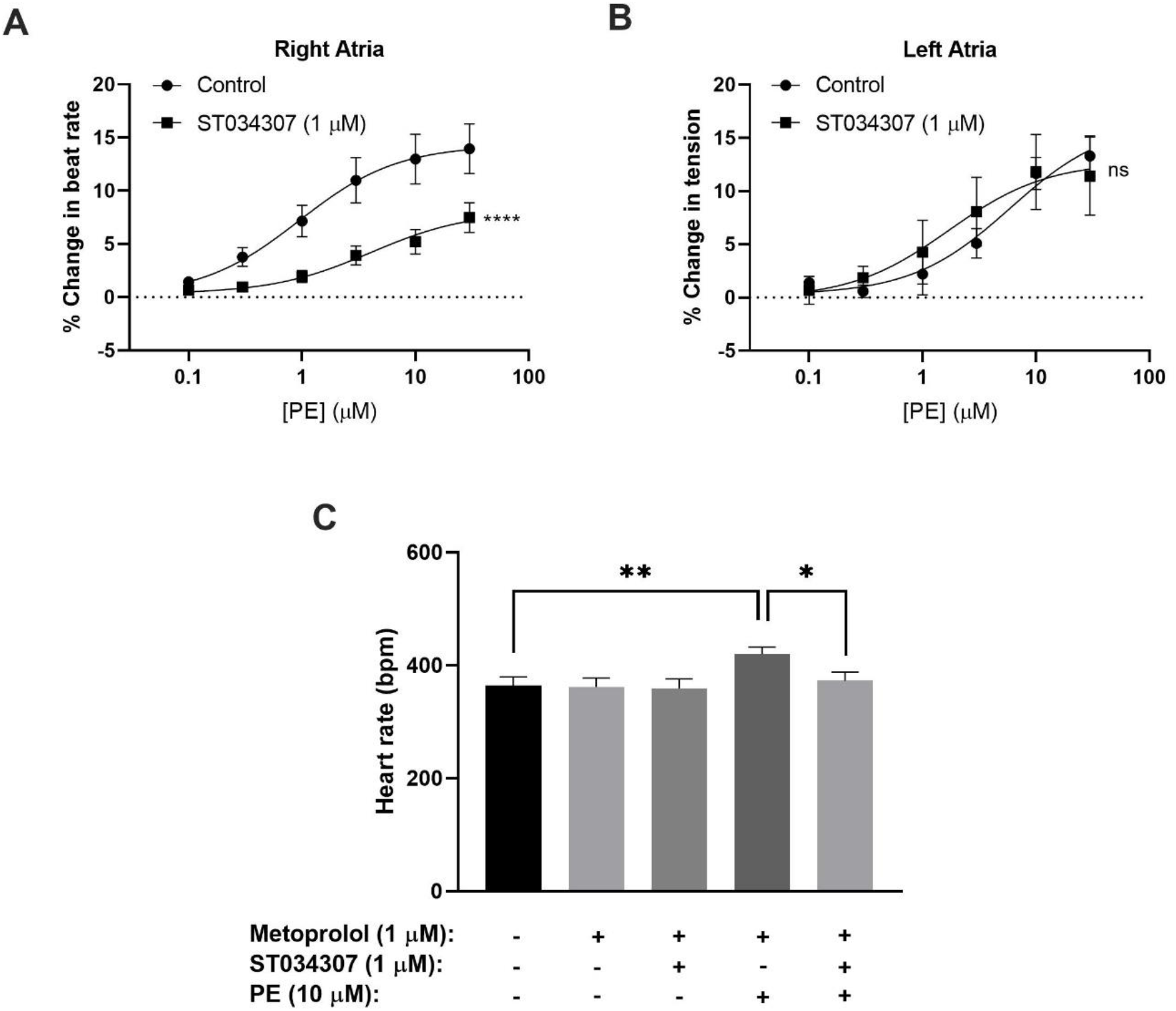
1 μM ST034307 inhibits changes in chronotropy, but not inotropy, in intact mouse right and left atria. **A**: Dose response curves to show the change in beating rate on cumulative addition of PE to spontaneously beating mouse right atria preparations under control conditions (circles, *n* = 15) and in the presence of 1 μM ST034307 (squares, *n* = 12). **B**: Dose response curves to show the change in tension generated on cumulative addition of PE to mouse left atria preparations under control conditions (circles, *n* = 5) and in the presence of 1 μM ST034307 (squares, *n* = 7). Dose-response curves in **A** and **B** (solid lines) were fitted using log(agonist) vs response (three-parameter model) using Graphpad Prism v9. Asterisks indicate significance level for effect of ST034307 compared to control at individual concentrations as determined using 2-way repeated measures ANOVA followed by Šídák’s multiple comparisons test. **C**: Bars represent mean heart rate (bpm) under different conditions as indicated below the x-axis. Significance bars represent comparison of the data using ANOVA followed by Fisher’s LSD test to compare all conditions (non-significant comparisons are not shown) (*n* = 11-14). Data are represented as mean ± SEM; *ns* = not significant; *, *P* < 0.05; **, *P* < 0.01; ***, *P* < 0.001.

Under control conditions, in the absence of ST034307 but in the presence of metoprolol, the spontaneous beating rate of right atria increased by a maximum of 14.46 % (95 % confidence interval (CI) = 12.23-17.05 %; *n* = 15) at 30 μM PE, with an EC50 of 0.91 μM (95 % CI = 0.38-2.2 μM) (Figure 2A, round symbols). 1 μM ST034307 reduced the response of beating rate to PE at all concentrations tested (Figure 2A, square symbols), with a maximum increase of 8.17 % (95 % CI = 6.12-12.60 %; *n* = 12) at 30 μM PE (EC50 = 3.86 μM; 95 % CI = 1.10-17.59 μM). This represented a significant overall reduction in the response to PE in the presence of ST034307 (*P* = 0.005, 2-way repeated measures ANOVA, PE vs. ST034307). For left atrial preparations (Figure 2B), PE increased the tension generated in response to electrical stimulation at 5 Hz. The maximum increase was 14.20 % (95 % CI = 11.24-18.53 %; *n* = 5) at 30 μM PE, with an EC50 of 4.38 μM (95 % CI = 1.92-18.53 μM). In the presence of ST034307, no difference was observed in the response to PE compared with control (*P* > 0.05, 2-way repeated measures ANOVA, PE vs. ST034307). The maximum change in tension generated in the presence of ST034307 was 16.32 % (95% CI = 11.73-23.34; *n* = 7) at 30 μM PE, with an EC50 of 0.91 μM (95 % CI = 0.82-10.82 μM). The basal heart rate of right atria after stabilisation and before addition of metoprolol was 364 ± 15.48 bpm (*n* = 12). No change in the right atrial basal heart rate was observed on addition of either metoprolol (362 ± 16.32 bpm) or metoprolol plus ST034307 (359 ± 16.79 bpm) before addition of PE, however the increase in beating rate induced by PE was inhibited in the presence of ST034307 as shown in Figure 2C.

### Inhibition of AC1 by ST034307 does not alter Ca^2+^ transient amplitude in isolated guinea pig atrial myocytes

To measure changes in cytosolic Ca^2+^ in response to PE, isolated guinea pig atrial myocytes were loaded with the cell-permeant Ca^2+^ sensitive dye Fluo-5F-AM. When stimulated at 1 Hz in PSS at 37 °C, guinea pig atrial myocytes exhibited the classical pattern of Ca^2+^ transient observed previously (Huser *et al*., 1996; Capel *et al*., 2021). We expressed calcium transient amplitude as change in mean cell fluorescence (F-F0)/F0 (Figures 3A-C). Addition of PE (10 μM) to the perfusion solution resulted in a 0.36-fold fold increase in Ca^2+^ transient amplitude from 3.56 ± 0.84 to 4.84 ± 1.04 (*P* = 0.009; *n* = 6; paired t-test) (Figures 3B and D). As shown in Figure 3E, addition of 1 μM ST034307 did not alter the basal Ca^2+^ transient amplitude compared to perfusion with PSS alone (PSS = 2.49 ± 0.33, *n* = 18; 1 μM ST = 1.94 ± 0.34, *n* = 15; P > 0.05, unpaired t-test). In the presence of 1 μM ST034307, a 0.39-fold increase in the Ca^2+^ transient amplitude was observed in response to 10 μM PE added to the perfusion solution, resulting in a Ca^2+^ transient amplitude of 2.69 ± 0.36 (*P* = 6.0×10^−4^; *n* = 15) (Figures 3C and F). Comparison of the percentage change in response to PE between control and 1 μM ST034307 confirmed that there was no significant difference between the amplitude of change in the presence of ST (*P* > 0.05, unpaired t-test).

**Figure 3.**
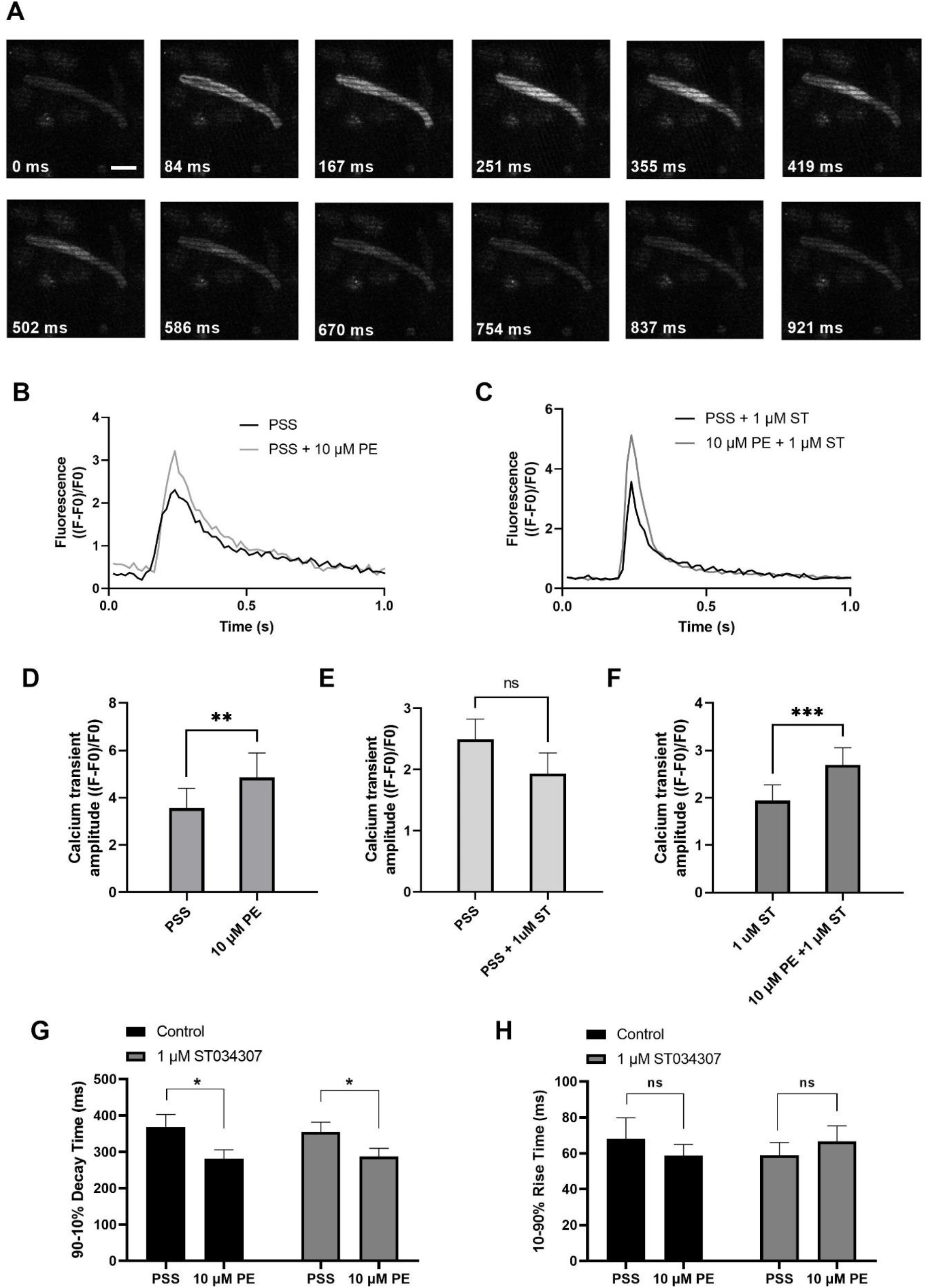
1 μM ST034307 does not alter the effect of PE on Ca^2+^ transients in isolated guinea pig atrial myocytes. **A**: Sequential images of a single atrial myocyte during the recording of a single Ca^2+^ transient, representing individual frames taken at the time points shown, representing time following start of the Ca^2+^ transient upstroke. Scale bar in first image = 20 μM. **B-C**: Representative average Ca^2+^ transients recoded from atrial myocytes during baseline recording (black trace) and following addition of PE (grey trace) in the absence (B) or presence (C) of 1μM ST034307. **D**: Effect of 10 μM PE on the Ca^2+^ transient amplitude recorded from isolated guinea pig atrial cells (*n* = 6; *N* = 3). **E**: Effect of 1 μM ST034307 on the basal Ca^2+^ transient amplitude before addition of PE (PE, *n* = 18; ST, *n* = 15; *N* = 3). **F**: Effect of 10 μM PE on the Ca^2+^ transient amplitude of cells in the presence of 1μM ST034307 (*n* = 15; *N* = 3). **G-H**: Effect of 10 μM PE on the 90-10% decay time (G) and 10-90% rise time (H) of Ca^2+^ transients recorded from cells perfused with PSS in the absence (black bars) or presence (grey bars) of 1 μM ST034307 before and after addition of 10 μM PE. All experiments were carried out at 35 ± 2°C and recordings were made 5 minutes following the start of each solution perfusion. Data are represented as mean ± SEM. Data in D and F were analysed using paired t-test. Data in E were analysed using unpaired t-test. Data in G and H were analysed using two-way, repeated measures ANOVA followed by Šídák’s multiple comparisons test; *ns* = not significant; *, *P* < 0.05; **, *P* < 0.01; ***, *P* < 0.001; *n* = cells; *N* = animals.

Further analysis of Ca^2+^ transient rise time and decay times confirmed that PE resulted in a significant decrease in the 90-10% decay time in control experiments from 369.48 ± 33.48 ms to 281.33 ± 24.78 ms (*P* = 0.02, 2-way ANOVA, *n* = 8) (Figure 3G, black bars) without changing the 10-90% rise time (Figure 3H, black bars). In the presence an of 1 μM ST034307, the same pattern was also observed with a decrease in 90-10% decay time from 355.00 ± 26.83 ms to 287.44 ± 22.47 ms (*P* = 0.01, 2-way ANOVA, *n* = 17) and no change in 10-90% rise time (Figures 3G and H, grey bars). These effects on decay and rise times were found not to differ between ST034307 and control experiments (*P* > 0.05, 2-way ANOVA).

### ST034307 inhibits SAN pacemaker activity in isolated adult guinea pig SAN cells

To assess the contribution of AC1 to rate and calcium signalling in SAN cells, the response to PE under control conditions and during AC1 inhibition with ST034307 was measured in isolated, spontaneously firing guinea pig SAN myocytes loaded with Fluo-5F-AM. Active spontaneously beating SAN cells were identified based on morphology as indicated by the black arrow in Figure 4A. Under control conditions, fluorophore-loaded SAN myocytes superfused with PSS at 35 °C spontaneously contracted at a rate of 103.59 ± 7.42 bpm (*n* = 5, Figure 4B, black bar). Although this rate is below the normal physiological rate for guinea pig SAN, lower rates are expected following the loading of cells with Fluo-5F-AM (Rigg & Terrar, 1996). Inclusion of 1 μM ST034307 in the perfusion solution resulted in a mean beating rate of 68.33 ± 5.07 bpm (*n* = 5, Figure 4B, grey bar) representing a significantly lower (0.34-fold) beat rate compared to control (*P* = 0.004, unpaired t-test). Superfusion of 10 μM PE led to a gradual increase in both calcium transient amplitude as well as the peak-to-peak firing rate over the course of 5 minutes as shown by the example traces in Figures 4C and D. On addition of 10 μM PE, the beat rate of SAN cells in the absence of ST034307 rose from 103.59 ± 7.42 bpm to a peak of 136.72 ± 8.56 bpm after 3 minutes, before decreasing to a plateau, corresponding to 117.37 ± 6.64 at 5 minutes (Figure 4E, *n* = 5). In the presence of ST034307, beat rate increased from 68.33 ± 5.07 bpm to 90.17 ± 9.663 after 60s, and remained unchanged for the remainder of the experiment, eg. 90.81 ± 8.7 bpm after 5 minutes (Figure 4E, *n* = 5). Overall, 1 μM ST034307 was found to significantly inhibit the rise in SAN cell spontaneous beat rate in response to superfusion with 10 μM PE (*P* = 0.02, 2-way repeated measures ANOVA). Qualitatively, this increase was no longer gradual but plateaued rapidly, being complete by the time of the 30 second recording (Figure 4E). Although qualitatively an increase in Ca^2+^ transient amplitude was observed in response to 10 μM PE, as shown by the representative traces in Figures 4C and D, quantitively this increase was not found to be significant, as shown in Figure 4F (*P* = 0.436, mixed effects analysis model). This is likely a consequence of the increased firing rate in response to PE, which would be expected to limit increases in calcium transient. However, in the presence of ST034307, mean Ca^2+^ transient amplitude was found to be consistently lower in the presence of ST034307 after 60s perfusion with PE (Figure 4F).

**Figure 4.**
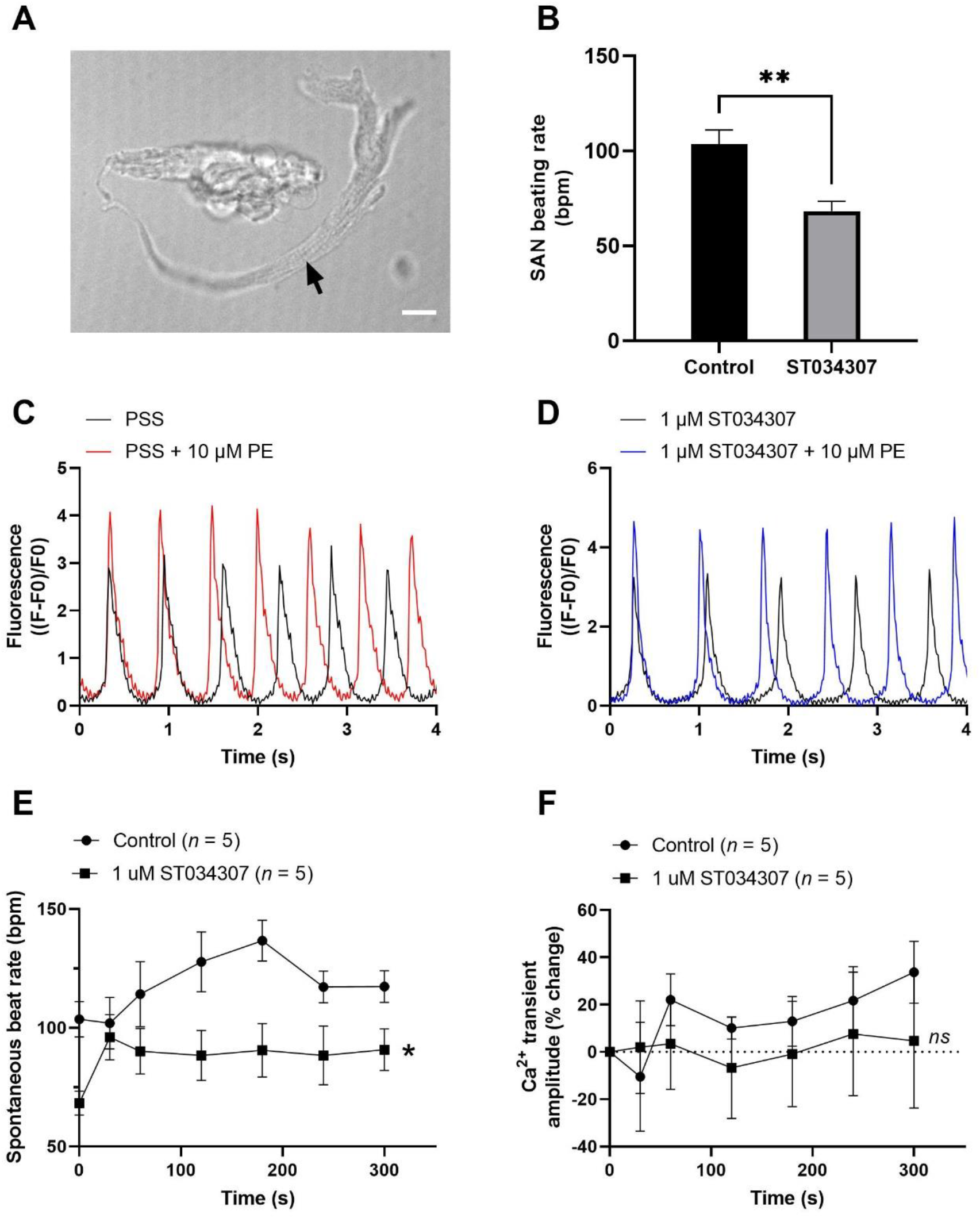
1 μM ST034307 inhibits the basal spontaneous beating rate and response to PE of isolated guinea pig SAN cells. **A**: Brightfield image to show example of an isolated guinea pig SAN cell as indicated by black arrow. Scale bar represents 10 μm. **B**: Effect of 1 μM ST034307 on the basal SAN cell Ca^2+^ transient amplitude before addition of PE (*n* = 5; *N* = 3-4). **C-D**: Representative 4 s recordings of changes in fluorescence (expressed as (F-F0)/F0) from spontaneously beating guinea pig SAN cells loaded with Fluo-5F-AM in PSS (C) or in PSS + 1 μM ST034307 (D). Black and red/blue traces represent recordings before and 5 minutes after addition of 10 μM PE respectively. Traces have been aligned according to the peak of the first transient. **E**: Effect of 10 μM PE on the spontaneous beating rate of isolated SAN cells in the presence (squares) an absence (circles) of 1 μM ST034307 (*n* = 5; *N* = 3-4). **F**: Effect of 10 μM PE on the Ca^2+^ transient amplitude recorded from isolated SAN cells in the presence (squares) an absence (circles) of 1 μM ST034307. Time in E and F represents time following addition of PE to the perfusion solution (*n* = 5; *N* = 3-4). All experiments were carried out at 35 ± 2°C. Data are represented as mean ± SEM. Data in B were analysed using unpaired t-test. Data in E were analysed using two-way, repeated measures ANOVA followed by Šídák’s multiple comparisons test. Data in F were analysed using mixed effects analysis followed by Šídák’s multiple comparisons test; *ns* = not significant; *, *P* < 0.05; **, *P* < 0.01; *n* = cells; *N* = animals.

## Discussion

Expression of IP_3_R2 in atrial cardiomyocytes is known to be six times greater in atrial myocytes compared to ventricular myocytes (Lipp *et al*., 2000), and expression of IP_3_R is known to be upregulated in human patients with chronic AF (Yamda *et al*., 2001) as well as the canine AF model (Zhao *et al*., 2007). The IP_3_ signalling pathway may therefore have potential as an atrial specific target for the treatment of AF (Tinker *et al*., 2016; Capel *et al*., 2021). There is increasing evidence that activation of the IP_3_ signalling pathway in atrial cells leads to the downstream activation of membrane bound Ca^2+^-sensitive adenylyl cyclases, principally AC1 and AC8 (Mattick *et al*., 2007; Burton & Terrar, 2021; Capel *et al*., 2021). Recently published work has demonstrated that inhibition of IP_3_R using 2-APB, and non-specific inhibition of ACs using MDL-12,330A, prevents the rise in spontaneous beating rate observed in response to PE in intact mouse right atria, as well as the rise in Ca^2+^ transient amplitude resulting from the intracellular release of caged-IP_3_ in isolated guinea pig atrial myocytes (Capel *et al*., 2021). The purpose of the current study was therefore to investigate the effect of pharmacological modulation of AC1, using the drug ST034307, to determine if AC1 may play a role in the downstream effects of IP_3_ signalling in cardiac tissue. Taken together, the data presented here suggest that AC1 plays a role in rate regulation at the sino-atrial node pacemaker but is less important in inotropic responses.

### The role of AC1 and IP_3_ signalling in the heart

Our immunohistochemistry work suggests that AC1 and IP_3_R colocalise in guinea-pig atrial myocytes, at least within the range of 0.16 μm, corresponding to the smallest pixel size in our immunocytochemistry images, (Figure 1). These data provide structural evidence in support of the hypothesis that IP_3_ mediated Ca^2+^ release in the cytosol could directly activate AC1 in atrial myocytes. Due to the relatively low resolution of fluorescent immunocytochemistry, the exact location of AC1, which may be dynamic, remains unclear. However, AC1 is known to be preferentially found in caveolae in rabbit SAN cells (Younes *et al*., 2008), while in mouse SAN cells, IP_3_R are known to be found in terminal SR in close proximity to the plasma membrane (Ju *et al*., 2011).

As shown in Figure 2A, 1μM ST034307 was found to significantly inhibit the increase in spontaneous beating rate of intact right atrial preparations by 56.5%. In contrast, ST034307 did not have a significant effect on the positive inotropic response to PE in paced murine left atria or potentiation of the Ca^2+^ transient in isolated guinea-pig atrial myocytes. There are several possible explanations for this observation. The simplest is that AC1 is minimally involved in the downstream effect of IP_3_ signalling in non-SAN atrial myocytes, but plays a more dominant role in the regulation of SAN pacemaker cells. Such an observation would be consistent with the observation that AC1 is preferentially expressed in the SAN and able to regulate I_f_ (Mattick *et al*., 2007). Alternatively, it is possible that the potentiation of AC2/5/6 by ST034307 hides the effects of AC1 inhibition on contractility in response to IP_3_ signalling in the atria. At higher concentrations (≥ 30 μM), ST034307 shows potentiation of AC2, and moderate potentiation of AC5/6, however these observations have not been reported at the lower concentration (1 μM) used in the present study (Brust *et al*., 2017). Indeed at 10 μM ST034307, we observed large variations in the response to PE, likely due to the contradictory effect of potentiation of these alternative ACs (data not shown).

Ca^2+^-sensitive ACs are implicated in atrial IP_3_ signalling since BAPTA, MDL-12,330A and W-7 (calmodulin inhibitor) all abolish the effects of PE on I_CaL_, (Wang et al., 2005). Moreover, as shown in Figure 1, AC1 is localised in close proximity with IP_3_Rs in atrial myocytes, meaning it is feasible that AC1 is activated directly by IP_3_-mediated Ca^2+^ release. It is possible however that IP_3_ mediated Ca^2+^ release can lead indirectly to AC1 activation via triggering other Ca^2+^ signals in different regions of the cells. One hypothesis that has been proposed is that local SR Ca^2+^ release from IP_3_R leads to a localized elevation in [Ca^2+^] at the ryanodine receptor, leading to amplification of RyR Ca^2+^ release and activation of L-type Ca^2+^ channels (LTCC) (Lipp *et al*., 2000; Liang *et al*., 2009). However, the abolition of response to exogenously applied IP_3_ in the presence of MDL appears to rule out this possibility, as this pathway would be expected to remain following inhibition of ACs (Capel *et al*., 2021). Another possibility however is that IP_3_R Ca^2+^ release triggers activation of AC1 indirectly via amplification of store operated Ca^2+^ entry (SOCE). In HEK293 cells, AC1 and AC8 are significantly activated by SOCE (Fagan *et al*., 1996) and SOCE is known to occur in close proximity to IP_3_Rs (Sampieri *et al*., 2018). Furthermore, it has been shown that AC8 interacts directly with Orai1, the pore forming subunit of SOCE channels (Willoughby *et al*., 2012). In addition, it has recently been demonstrated that in HeLa cells, activation of IP_3_R clusters tethered below the plasma membrane by the KRas-induced actin-interacting protein (KRAP) leads to localised depletion of ER Ca^2+^, which in turn leads to SOCE via the activation of stromal interaction molecule 1 (STIM1) (Thillaiappan *et al*., 2021). Interestingly, in isolated mouse SAN cells, the SOCE inhibitor SKF-9665 inhibited Ca^2+^ influx in SAN in response to pharmacological SR unloading and reduced the spontaneous rate by 27% in these conditions (Ju *et al*., 2007). It remains to be explored whether this finding involves Ca^2+^ -activated adenylyl cyclases. The role of IP_3_ signalling in activation of SOCE in cardiac cells, and the potential for downstream regulation of AC activity therefore warrants future investigation.

### The role of AC1 and IP_3_ signalling in cardiac pacemaker activity

Pacemaker activity in mouse SAN cells can undergo modulation by both IP_3_ agonists and antagonists, and this modulation can be abolished following IP_3_R2 knock-out, demonstrating that IP_3_ signalling can play a role in regulating pacemaker activity (Ju *et al*. 2011). In addition, IP_3_ has been shown to induce Ca^2+^ sparks in close proximity to the surface membrane in pacemaker cells, and it has been suggested that this may lead to modulation of inward Na^+^/Ca^2+^ exchange current or activation of alternative Ca^2+^ dependent currents (Ju *et al*., 2012). Both IP_3_R2 (Ju *et al*. 2011) and AC1 (Mattick *et al*. 2007; Younes *et al*., 2008) are expressed in the SAN, and there is therefore the potential for activation of Ca^2+^-sensitive adenylyl cyclases downstream of IP_3_R Ca^2+^ release in SAN cells as well as in non-pacemaker cells (Mattick *et al*. 2007; Ju *et al*. 2012). These observations support a role for the Ca^2+^-activated adenylyl cyclases AC1 and AC8 in atrial and SAN IP_3_ signalling, however the involvement of other adenylyl cyclases cannot be ruled out based on these data alone.

1 μM ST034307 significantly reduced the positive chronotropic effect of PE in intact mouse right atria (Figure 2A) without altering inotropy in the left atria (Figure 2B). Similarly, 1 μM ST034307 reduced spontaneous Ca^2+^ transient firing rate in isolated guinea-pig SAN cells (Figure 4B), but did not inhibit increases in Ca^2+^ transient amplitude in response to PE in either atrial (Figure 3) or SAN cells (Figure 4). Although it has been shown that ST034307 can potentiate AC2, AC5 and AC6 activity at higher concentrations over 30 μM, (Brust *et al*., 2017) the net effect of ST034307 in the SAN is expected to favour the effect of AC1 inhibition due to the higher concentration (Mattick *et al*., 2007; Younes *et al*., 2008) and activity of AC1 (Younes *et al*., 2008) compared to AC2, AC5 and AC6 within SAN cells. Furthermore, there is a reduction in the maximal responses to PE but the EC_50_ is unchanged (Figure 2A), suggesting the effects of ST034307 are via inhibition of a target in the IP_3_ signalling pathway. The most likely explanation for these observations is therefore that AC1 is activated downstream of IP_3_-mediated Ca^2+^ release and that AC1 inhibition by ST034307 thus inhibits the downstream effectors of IP_3_ signalling in the SAN. Pertinently, AC1 is implicated as being directly involved in the positive chronotropic effect of the IP_3_ signalling pathway since application of the non-specific AC inhibitor MDL-12,330A abolishes the positive chronotropic response to PE in the absence of β-adrenergic signalling (Capel *et al*., 2021), thus supporting that cAMP production by ACs is involved in the chronotropic response to IP_3_R activation. Consistent with this hypothesis, AC1 and AC8 are present in the SAN plasma membrane and either or both isoforms potentiate the pacemaker current (Mattick *et al*., 2007). Furthermore, both AC1 (Younes *et al*., 2008) and IP_3_R2 localise in close proximity to caveolae in SAN cells (Barbuti *et al*., 2004; Ju *et al*., 2011), and the immunocytochemistry data reported in the current paper together with that reported in Capel *et al*. (2021) in atrial cells provide further support for the direct activation of AC1 by IP_3_-mediated Ca^2+^ release.

1 μM ST034307 did not abolish the response to PE in right atrial tissue (Figure 2A), although based on the reported IC_50_ of 2.3 μM (Brust *et al*., 2017), this dose of ST may have been insufficient to cause maximal AC1 inhibition. At higher doses however contradictory results were observed, likely due to the potentiation of alternative AC isoforms as discussed below and as reported by Brust *et al*. (2017). Whilst our findings suggest a role for AC1 downstream of IP_3_ mediated Ca^2+^ release, in the absence of a specific AC8 inhibitor, our results cannot rule out the possibility that AC8 is also involved. At the time of writing no such inhibitor is commercially available.

The specificity of cAMP signalling is known to rely on localisation within micro-(Zaccolo & Pozzan, 2002) and nanodomains (Surdo *et al*., 2017). We therefore hypothesise that the same is true for cAMP activated downstream to IP_3_ signalling, and such localisation would explain why the specificity of this Ca^2+^ signal is not lost despite constant global Ca^2+^ transients within SAN cells. Moreover, since AC1 regulation by Ca^2+^ is biphasic (Fagan *et al*., 1996), it is possible that IP_3_ induced stimulation of AC1 only occurs after cytosolic Ca^2+^ and the membrane potential have decreased, meaning this additional cAMP is likely only produced during the early and late phase of diastole allowing stimulation of I_f_ by AC1 at the correct time point. Our findings suggest that within SAN cells, Ca^2+^ released from IP_3_R can activate either directly or indirectly AC1 and that this can modulate both basal and stimulated pacemaking mechanisms. Such a mechanism would be comparable but independent of that by which the release of Ca^2+^ from RyR (Rigg & Terrar, 1996; Bogdanov *et al*., 2001), or Ca^2+^ influx via the L-type (Mangoni *et al*., 2003; Jones *et al*., 2007) or T-type Ca^2+^ channels (Huser *et al*., 1996) regulates basal pacemaking.

### Clinical relevance

Abnormal Ca^2+^ signalling underlies the pathology of many forms of cardiac disorders and arrythmias, including AF (Landstrom *et al*., 2017). Current rate control medication for diseases such as heart failure and atrial fibrillation target either β-adrenergic signalling, Na^+^/K^+^-ATPase, or I_f_, while few selective pharmacological treatments exist for sinus node dysfunction. Our findings that IP_3_ triggered AC1 activity and thus the cAMP signals it produces regulate pacemaking may provide a potential new avenue for novel pharmacological treatments via the modulation of this IP_3_-Ca^2+^-AC1-cAMP pathway. Sinus node dysfunction (or sick sinus syndrome) comprises a group of progressive non-curable diseases where the heart rate is inappropriately bradycardic or tachycardic, resulting in increased morbidity rates (Alonso *et al*., 2014). In familial sinus node dysfunction, multiple different mutated proteins have been implicated including key Ca^2+^ handling proteins such as RyR2, calsequestrin and Cav1.3, alongside HCN4 (Wallace *et al*., 2021). The crucial importance of Ca^2+^ handling and signalling to pacemaking and apparent importance in sinus node dysfunction makes this an important potential target for the modulation of pacemaking in patients with sinus node dysfunction as well as patients with heart failure and atrial fibrillation. However, a directed approach to finely modulate SAN Ca^2+^ signalling directly is as yet to be described.

### Limitations of this study

ST034307 is known to be non-specific, potentiating AC2, AC5 and AC6 activity at concentrations of 30 μM or over (Brust et al, 2017). Indeed, in our preliminary experiments we found responses to ST034307 to be highly variable and unstable at 10 μM (data not shown). As such we used 1 μM ST034307 in our experiments, a concentration below the reported IC_50_ of 2.3 μM (Brust *et al*., 2017). Chromones have a widely documented biological activity in a range of biological settings (Gaspar *et al*., 2014), and as such, the possibility of off target effects of this drug cannot be eliminated. Further investigations using the genetic knockdown or knockout of AC1 in cardiac tissue, including the SAN, may therefore provide a more comprehensive understanding of the role played by AC1 downstream of IP_3_ signalling in the heart. Similarly, as ST034307 has a bicyclic planar structure and is derived from the compound NB001 that has an adenine motif (Brust *et al*., 2017), it is possible that the mechanism of action involves binding to ATP binding cassettes in adenylyl cyclases. This raises the possibility of ST034307 affecting multiple different targets containing nucleotide-binding cassettes.

In the present study, we chose to investigate this pathway in mouse whole atria and guinea pig isolated cells. Guinea pig cells were chosen specifically due to the similarities between human and guinea pig cardiac electrophysiology, however future studies using human derived tissue will be required to fully understand the role of cardiac AC1 in humans.

### Conclusion

The present study highlights a role for the Ca^2+^-dependent AC isoform AC1 in determining cardiac pacemaking, both at the level of the isolated SAN cell as well as at the level of the intact beating right atria. Due to the question relating to non-specificity of ST034307, and the lack of more specific compounds targeting AC1, definitive demonstration of the involvement of AC1 in SAN IP_3_ signalling or evaluation of the role of AC1 in atrial myocyte IP_3_ signalling is complex based solely on these experiments. This work therefore highlights the need for the development of specific AC1 inhibitors that would overcome issues of compensation associated with AC1 single knock-out in genetically modified mouse strains. Regardless, the most likely explanation for the blunting of the positive chronotropic response of the SAN to PE, in this study, is inhibition of AC1 by ST034307. Moreover, the cause of the divergent effects of 1μM ST034307 between the SAN and atrial myocytes merits further investigation of the mechanisms regulating both atrial and SAN IP_3_ signalling.

Overall, this study supports the existence of an IP_3_ → AC1 → cAMP signalling pathway regulating SAN pacemaking in response to α-adrenergic signalling. The findings presented support a role for AC1 downstream of IP_3_-mediated Ca^2+^ release, providing a new example of how crosstalk between Ca^2+^ and cAMP signalling is involved in regulating SAN pacemaker activity. These data add to the already published mechanism of cross-talk between Ca^2+^ and cAMP signalling within the SAN, with Ca^2+^ already having been shown to control basal cAMP levels and pacemaker activity (Mattick *et al*., 2007; Younes *et al*., 2008), and provide further support for previous work identifying a link between IP_3_ signalling and the downstream activation of Ca^2+^-dependent ACs (Capel *et al*., 2021). Furthermore, the concept that SAN IP_3_ signalling and automaticity can be targeted through cyclic nucleotide signalling suggests further investigation of putative IP_3_-cAMP signalling pathways in cardiac atria may identify novel targets, for example phosphodiesterases, to modulate pacemaking and also prevent the triggering of atrial arrhythmias such as atrial fibrillation.

## Author Contributions Statement

R.A.B.B. conceived the research. DT, MZ and AR contributed intellectually to the study. SJB and RABB designed the study. SJB and MR carried out intact atrial preparations. SJB, MR and RC carried out SAN isolated cell work. EA and TA carried out immunofluorescence work and produced Figure 1. SJB and MR wrote the manuscript. SJB, RAC an EA carried out animal dissections and cell isolations. All authors have contributed to refinement of the manuscript.

## Acknowledgements

We acknowledge the support of Dr Tim Viney and Dr Thomas P. Collins, Department of Pharmacology, University of Oxford for their assistance with this manuscript. SJB is a post-doctoral scientist funded by the British Heart Foundation (PG/18/4/33521). RABB is funded by a Sir Henry Dale Wellcome Trust and Royal Society Fellowship (109371/Z/15/Z) and holds a Senior Research Fellowship at Linacre College, Oxford. RAC is a post-doctoral scientist funded by the Wellcome Trust and Royal Society (109371/Z/15/Z). MZ is supported by the British Heart Foundation (RG/17/6/32944). TA received funding from the Returners Carers Fund (PI RABB), Medical Science Division, University of Oxford, the Nuffield Benefaction for Medicine and the Wellcome Institutional Strategic Support Fund (ISSF), University of Oxford. EA received funding from the Returners Carers Fund (PI RAC), University of Oxford. R.A.B.B. and M.Z. acknowledge support from the BHF Centre of Research Excellence, Oxford. R.A.B.B acknowledges support from the Covid-19 Rebuilding Research Momentum Fund (CRRMF) Oxford Funds.

## Figure Legends

**Supplementary video (related to Figure 1)**

Supplementary video shows representative z-stack images from fixed, isolated guinea pig atrial myocytes immunolabelled for IP_3_R2 (magenta) and AC1 (cyan). Z-stacks were recorded at 1 μm intervals.

## References

Alonso A, Jensen PN, Lopez FL, Chen LY, Psaty BM, Folsom AR & Heckbert SR. (2014). Association of Sick Sinus Syndrome with Incident Cardiovascular Disease and Mortality: The Atherosclerosis Risk in Communities Study and Cardiovascular Health Study. Plos One 9.

Ando H, Mizutani A, Kiefer H, Tsuzurugi D, Michikawa T & Mikoshiba K. (2006). IRBIT suppresses IP_3_ receptor activity by competing with IP3 for the common binding site on the IP3 receptor. Molecular Cell 22, 795–806.

Barbuti A, Gravante B, Riolfo M, Milanesi R, Terragni B & DiFrancesco D. (2004). Localization of pacemaker channels in lipid rafts regulates channel kinetics. Circulation Research 94, 1325–1331.

Bers DM. (2002). Cardiac excitation-contraction coupling. Nature 415, 198–205.

Bogdanov KY, Vinogradova TM & Lakatta EG. (2001). Sinoatrial nodal cell ryanodine receptor and Na^+^-Ca^2+^ exchanger - Molecular partners in pacemaker regulation. Circulation Research 88, 1254–1258.

Brand T. (2005). The Popeye domain-containing gene family. Cell Biochem Biophys 43, 95– 103.

Brust TF, Alongkronrusmee D, Soto-Velasquez M, Baldwin TA, Ye ZS, Dai MJ, Dessauer CW, van Rijn RM & Watts VJ. (2017). Identification of a selective small-molecule inhibitor of type 1 adenylyl cyclase activity with analgesic properties. Science Signaling 10.

Burton R-AB & Terrar DA. (2021). Emerging Evidence for cAMP-calcium Cross Talk in Heart Atrial Nanodomains Where IP_3_-Evoked Calcium Release Stimulates Adenylyl Cyclases. Contact 4, 1–13.

Capel RA, Bose SJ, Collins TP, Rajasundaram S, Ayagama T, Zaccolo M, Burton RAB & Terrar DA. (2021). IP_3_-mediated Ca^2+^ release regulates atrial Ca^2+^ transients and pacemaker function by stimulation of adenylyl cyclases. American Journal of Physiology-Heart and Circulatory Physiology 320, H95–H107.

Capel RA & Terrar DA. (2015). The importance of Ca(2+)-dependent mechanisms for the initiation of the heartbeat. Front Physiol Mar 25, 6:80.

Chen-Izu Y, Xiao RP, Izu LT, Cheng H, Kuschel M, Spurgeon H & Lakatta EG. (2000). Gi-dependent localization of beta2-adrenergic receptor signaling to L-type Ca^2+^ channels. Biophys J 79, 2547–2556.

Collins TP, Bayliss R, Churchill GC, Galione A & Terrar DA. (2011). NAADP influences excitation-contraction coupling by releasing calcium from lysosomes in atrial myocytes. Cell calcium 50, 449–458.

Collins TP & Terrar DA. (2012). Ca(2+)-stimulated adenylyl cyclases regulate the L-type Ca(2+) current in guinea-pig atrial myocytes. J Physiol 590, 1881–1893.

de Rooij J, Zwartkruis FJ, Verheijen MH, Cool RH, Nijman SM, Wittinghofer A, & Bos JL. (1998). Epac is a Rap1 guanine-nucleotide-exchange factor directly activated by cyclic AMP. Nature 396, 474–477.

Difrancesco D, Noble D & Denyer JC. (1991). The contribution of the pacemaker current (I_f_) to generation of spontaneous activity in rabbit sinoatrial node myocytes. Journal of Physiology-London 434, 23–40.

Difrancesco D & Tortora P. (1991). Direct activation of cardiac-pacemaker channels by intracellular cyclic-AMP. Nature 351, 145–147.

Difrancesco D & Tromba C. (1988). Muscarinic control of the hyperpolarization-activated current (I_f_) in rabbit sino-atrial node myocytes. Journal of Physiology-London 405, 493–510.

Domeier TL, Zima AV, Maxwell JT, Huke S, Mignery GA & Blatter LA. (2008). IP3 receptor-dependent Ca^2+^ release modulates excitation-contraction coupling in rabbit ventricular myocytes. Am J Physiol Heart Circ Physiol 294, H596–604.

Fagan KA, Mahey R & Cooper DMF. (1996). Functional co-localization of transfected Ca2+-stimulable adenylyl cyclases with capacitative Ca2+ entry sites. Journal of Biological Chemistry 271, 12438–12444.

Fesenko EE, Kolesnikov SS, & Lyubarsky AL. (1985). Induction by cyclic GMP of cationic conductance in plasma membrane of retinal rod outer segment. Nature 313, 310– 313.

Gaspar A, Matos MJ, Garrido J, Uriarte E & Borges F. (2014). Chromone: A Valid Scaffold in Medicinal Chemistry. Chemical Reviews 114, 4960–4992.

Georget M, Mateo P, Vandecasteele G, Jurevicius J, Lipskaia L, Defer, Hanoune J, Hoerter J & Fischmeister R (2002). Augmentation of cardiac contractiliy with no change in L-type Ca^2+^ current in transgenic mice with a cardiac-directed expression of the human adenylyl cyclase type 8 (AC8). FASEB J 16(12), 1636–1638.

Hancox JC & Mitcheson JS. (1997). Ion channel and exchange currents in single myocytes isolated from the rabbit atrioventricular node. Canadian Journal of Cardiology 13, 1175–1182.

Harvey RD & Clancy CE. (2021). Mechanisms of cAMP compartmentation in cardiac myocytes: experimental and computational approaches to understanding. J Physiol 599, 4527–4544.

Hattori M, Suzuki AZ, Higo T, Miyauchi H, Michikawa T, Nakamura T, Inoue T & Mikoshiba K. (2004). Distinct roles of inositol 1,4,5-trisphosphate receptor types 1 and 3 in Ca2+ signaling. Journal of Biological Chemistry 279, 11967–11975.

Huser J, Lipsius SL & Blatter LA. (1996). Calcium gradients during excitation-contraction coupling in cat atrial myocytes. Journal of Physiology-London 494, 641–651.

Jones SA, Boyett MR & Lancaster MK. (2007). Declining into failure -The age-dependent loss of the L-type calcium channel within the sinoatrial node. Circulation 115, 1183–1190.

Ju YK, Chu Y, Chaulet H, Lai D, Gervasio OL, Graham RM, Cannell MB & Allen DG. (2007). Store-operated Ca2+ influx and expression of TRPC genes in mouse sinoatrial node. Circulation Research 100, 1605–1614.

Ju YK, Liu J, Lee BH, Lai D, Woodcock EA, Lei M, Cannell MB & Allen DG. (2011). Distribution and functional role of inositol 1,4,5-trisphosphate receptors in mouse sinoatrial node. Circ Res 109, 848–857.

Ju YK, Woodcock EA, Allen DG & Cannell MB. (2012). Inositol 1,4,5-trisphosphate receptors and pacemaker rhythms. Cell. Cardiol. 53, 375–381.

Katsushika S, Chen L, Kawabe JI, Nilakantan R, Halnon NJ, Homcy CJ & Ishikawa Y. (1992). Cloning and characterization of a 6th adenylyl cyclase isoform - type-V and type-VI constitute a subgroup within the mammalian adenylyl cyclase family. Proceedings of the National Academy of Sciences of the United States of America 89, 8774–8778.

Krebs EG, & Beavo JA, (1979). Phosphorylation-dephosphorylation of enzymes. Annu Rev Biochem 48, 923–959.

Lakatta EG, Maltsev VA & Vinogradova TM. (2010). A coupled SYSTEM of intracellular Ca2+ clocks and surface membrane voltage clocks controls the timekeeping mechanism of the heart’s pacemaker. Circ Res 106, 659–673.

Landstrom AP, Dobrev D & Wehrens XHT. (2017). Calcium Signaling and Cardiac Arrhythmias. Circulation Research 120, 1969–1993.

Liang X, Xie H, Zhu PH, Hu J, Zhao Q, Wang CS & Yang C. (2009). Enhanced activity of inositol-1,4,5-trisphosphate receptors in atrial myocytes of atrial fibrillation patients. Cardiology 114, 180–191.

Lipp P, Laine M, Tovey SC, Burrell KM, Berridge MJ, Li WH & Bootman MD. (2000). Functional InsP(3) receptors that may modulate excitation-contraction coupling in the heart. Current Biology 10, 939–942.

Lou LX, Li CH, Wang J, Wu AM, Zhang T, Ma Z, Chai LM, Zhang DM, Zhao YZ, Nie B, Jin QS, Chen HY & Liu WJ. (2021). Yiqi Huoxue preserves heart function by upregulating the Sigma-1 receptor in rats with myocardial infarction. Experimental and Therapeutic Medicine 22.

Mangoni ME, Couette B, Bourinet E, Platzer J, Reimer D, Striessnig J & Nargeot J. (2003). Functional role of L-type Ca(v)13Ca(2+) channels in cardiac pacemaker activity. Proceedings of the National Academy of Sciences of the United States of America 100, 5543–5548.

Mattick P, Parrington J, Odia E, Simpson A, Collins T & Terrar D. (2007). Ca2+-stimulated adenylyl cyclase isoform AC1 is preferentially expressed in guinea-pig sino-atrial node cells and modulates the I(f) pacemaker current. J Physiol 582, 1195–1203.

Premont RT, Chen JQ, Ma HW, Ponnapalli M & Iyengar R. (1992). 2 members of a widely expressed subfamily of hormone-stimulated adenylyl cyclases. Proceedings of the National Academy of Sciences of the United States of America 89, 9809–9813.

Rigg L, Mattick PAD, Heath BM & Terrar DA. (2003). Modulation of the hyperpolarization-activated current (I-f) by calcium and calmodulin in the guinea-pig sino-atrial node. Cardiovascular Research 57, 497–504.

Rigg L & Terrar DA. (1996). Possible role of calcium release from the sarcoplasmic reticulum in pacemaking in guinea-pig sino-atrial node. Experimental Physiology 81, 877–880.

Salvador JBI & Egger M. (2018). Obstruction of ventricular Ca2+-dependent arrhythmogenicity by inositol 1,4,5-trisphosphate-triggered sarcoplasmic reticulum Ca2+ release. Journal of Physiology-London 596, 4323–4340.

Sampieri A, Santoyo K, Asanov A & Vaca L. (2018). Association of the IP3R to STIM1 provides a reduced intraluminal calcium microenvironment, resulting in enhanced store-operated calcium entry. Scientific Reports 8.

Surdo NC, Berrera M, Koschinski A, Brescia M, Machado MR, Carr C, Wright P, Gorelik J, Morotti S, Grandi E, Bers DM, Pantano S & Zaccolo M. (2017). FRET biosensor uncovers cAMP nano-domains at beta-adrenergic targets that dictate precise tuning of cardiac contractility. Nature Communications 8.

Terrar DA. (2020). Calcium Signaling in the Heart. Calcium Signaling, 2nd Edition 1131, 395–443.

Thillaiappan NB, Smith HA, Atakpa-Adaji P & Taylor CW. (2021). KRAP tethers IP3 receptors to actin and licenses them to evoke cytosolic Ca2+ signals. Nature Communications 12.

Tinker A, Finlay M, Nobles M & Opel A. (2016). The contribution of pathways initiated via the G(q\11) G-protein family to atrial fibrillation. Pharmacological Research 105, 54–61.

Tsutsui K, Monfredi OJ, Sirenko-Tagirova SG, Maltseva LA, Bychkov R, Kim MS, Ziman BD, Tarasov KV, Tarasova YS, Zhang J, Wang MY, Maltsev AV, Brennan JA, Efimov IR, Stern MD, Maltsev VA & Lakatta EG. (2018). A coupled-clock system drives the automaticity of human sinoatrial nodal pacemaker cells. Science Signaling 11.

Uchida K, Aramaki M, Nakazawa M, Yamagishi C, Makino S, Fukuda K, Nakamura T, Takahashi T, Mikoshiba K & Yamagishi H. (2010). Gene Knock-Outs of Inositol 1,4,5-Trisphosphate Receptors Types 1 and 2 Result in Perturbation of Cardiogenesis. Plos One 5.

Vinogradova TM, Sirenko S, Lyashkov AE, Younes A, Li Y, Zhu W, Yang D, Ruknudin AM, Spurgeon H & Lakatta EG. (2008). Constitutive phosphodiesterase activity restricts spontaneous beating rate of cardiac pacemaker cells by suppressing local Ca2+ releases. Circ Res 102, 761–769.

Vinogradova TM, Zhou Y, Bogdanov KY, Yang D, Kuschel M, Cheng H & Xiao RP. (2000). Sinoatrial node pacemaker activity requires Ca^2+^/Calmodulin-dependent kinase II activation. Circ Res 87, 760–767.

Wallace MJ, El Refaey M, Mesirca P, Hund TJ, Mangoni ME & Mohler PJ. (2021). Genetic Complexity of Sinoatrial Node Dysfunction. Frontiers in Genetics 12.

Walsh DA, Perkins JP, & Krebs EG. (1968) An adenosine 3’,5’-monophosphate dependant protein kinase from rabbit skeletal muscle. J Biol Chem 243, 3763–3765.

Willoughby D, Everett KL, Halls ML, Pacheco J, Skroblin P, Vaca L, Klussmann E & Cooper DMF. (2012) Direct binding between Orai1 and AC8 mediates dynamic interplay between Ca^2+^ and cAMP signaling. Science Signaling 5(219), ra29

Yamda J, Ohkusa T, Nao T, Ueyama T, Yano M, Kobayashi S, Hamano K, Esato K & Matsuzaki M. (2001). Up-regulation of inositol 1,4,5 trisphosphate receptor expression in atrial tissue in patients with chronic atrial fibrillation. J Am Coll Cardiol 37, 1111–1119.

Yaniv Y, Spurgeon HA, Ziman BD, Lakatta EG. (2013). Ca^2+^/Calmodulin-Dependent Protein Kinase II (CaMKII) Activity and Sinoatrial Nodal Pacemaker Cell Energetics. PLoS One 8, e57079.

Younes A, Lyashkov AE, Graham D, Sheydina A, Volkova MV, Mitsak M, Vinogradova TM, Lukyanenko YO, Li Y, Ruknudin AM, Boheler KR, van Eyk J & Lakatta EG. (2008). Ca(2+) -stimulated basal adenylyl cyclase activity localization in membrane lipid microdomains of cardiac sinoatrial nodal pacemaker cells. J Biol Chem 283, 14461–14468.

Zaccolo M & Pozzan T. (2002). Discrete microdomains with high concentration of cAMP in stimulated rat neonatal cardiac myocytes. Science 295, 1711–1715.

Zaccolo M, Zerio A & Lobo MJ. (2021) Subcellular organisation of the cAMP signalling pathway. Pharm Rev 73, 278–309.

Zhao ZH, Zhang HC, Xu Y, Zhang P, Li XB, Liu YS & Guo JH. (2007). Inositol-1,4,5-trisphosphate and ryanodine-dependent Ca^2+^ signaling in a chronic dog model of atrial fibrillation. Cardiology 107, 269–276.

